# Modelling the transcription factor DNA-binding affinity using genome-wide ChIP-based data

**DOI:** 10.1101/061978

**Authors:** Monther Alhamdoosh, Dianhui Wang

## Abstract

Understanding protein-DNA binding affinity is still a mystery for many transcription factors (TFs). Although several approaches have been proposed in the literature to model the DNA-binding specificity of TFs, they still have some limitations. Most of the methods require a cut-off threshold in order to classify a K-mer as a binding site (BS) and finding such a threshold is usually done by handcraft rather than a science. Some other approaches use a prior knowledge on the biological context of regulatory elements in the genome along with machine learning algorithms to build classifier models for TFBSs. Noticeably, these methods deliberately select the training and testing datasets so that they are very separable. Hence, the current methods do not actually capture the TF-DNA binding relationship. In this paper, we present a threshold-free framework based on a novel ensemble learning algorithm in order to locate TFBSs in DNA sequences. Our proposed approach creates TF-specific classifier models using genome-wide DNA-binding experiments and a prior biological knowledge on DNA sequences and TF binding preferences. Systematic background filtering algorithms are utilized to remove non-functional K-mers from training and testing datasets. To reduce the complexity of classifier models, a fast feature selection algorithm is employed. Finally, the created classifier models are used to scan new DNA sequences and identify potential binding sites. The analysis results show that our proposed approach is able to identify novel binding sites in the Saccharomyces cerevisiae genome.

**Availability:** http://homepage.cs.latrobe.edu.au/dwang/DNNESCANweb

## 1 Introduction

Deciphering the regulation mechanism of gene expression is an important stage towards understanding how cells manipulate nonprotein-coding DNA sequences. The binding of transcription factor (TF) proteins upstream, downstream or on the introns of genes is one of the most challenging problems to be investigated in bioinformatics. A TF protein binds on short DNA elements, called transcription factor binding sites (TFBSs), so that it down regulates or up regulates the expression of associated genes (Stormo, 2010). Although the advancement of *in vivo* experimental technologies of DNA-protein interaction, e.g., protein binding microarrays (Bulyk, 2007), has enabled researchers to identify TFBSs effortlessly, the accurate *in silico* prediction of TFBSs helps us better understand TF protein-DNA binding mechanisms. However, the difficulty of locating TF-BSs using computational approaches emerges for two main reasons: (i) TFBSs are short and degenerate (5-20 bp) (Stormo, 2000), and (ii) the specificity of protein-DNA binding does not depend only on DNA bases (base readout), but it also depends on the 3D structures of DNA and TF protein macromolecules (shape readout) (Rohs *et al.*, 2010).

Several technical limitations should be seriously considered when a computational system is developed for the identification of binding sites. First, most of the current approaches use a cut-off threshold to decide whether a K-mer (DNA sequence of length K) is a binding site or a background sequence. The selection of an appropriate threshold is not straightforward. Too small a threshold increases the number of false positives (low precision) while too large a threshold removes many true positives (low recall). Therefore, a threshold-free system to locate binding sites is highly recommended (Wang *et al.*, 2016). Second, the majority of computational methods in the literature use very simple models to represent binding sites, i.e., template-based strategies. They do not utilize the complex relationships between different DNA and TF protein characteristics (Gunewardena *et al.*, 2006; Burden and Weng, 2005; Oshchepkov *et al.*, 2004; Liu *et al.*, 2001; Ponomarenko *et al.*, 1999; Karas *et al.*, 1996). Hence, efficient models that can learn these relationships would be extremely useful in order to understand the different mechanisms that control the binding of TF proteins to DNA regulatory elements (Wang *et al.*, 2016; Hooghe *et al.*, 2012). Third, the current machine learning-based approaches construct the learner models using known binding sites and *randomly* selected background K-mers from the same genome (Hooghe *et al.*, 2012; Bauer *et al.*, 2010; Meysman *et al.*, 2011; Steffen *et al.*, 2002; Pudimat *et al.*, 2005). Such training sequence-sets are more likely to be quite separable. As a result, the TFBS classifier might not be able to recognize background K-mers that have the same nucleotide sequence of a true binding site. To overcome this limitation and accurately model TFBSs, the classifier models should have the ability to learn decision boundaries from a wide range of DNA sequences.

The computational approaches for locating TFBSs can be grouped into two main categories (Hooghe *et al.*, 2012): sequence-dependent approaches that identify binding sites based on the DNA bases of a given sequence (Gunewardena *et al.*, 2006; Bi *et al.*, 2011; Broos *et al.*, 2011; Wang *et al.*, 2016; Hooghe *et al.*, 2012; Bauer *et al.*, 2010) and structure-dependent approaches that predict the interactions between binding site DNA bases and TF protein amino acids by analyzing the resolved 3D structures of TF protein-DNA complexes (Piovesan *et al.*, 2012; Angarica *et al.*, 2008; Jamal Rahi *et al.*, 2008; Kaplan *et al.*, 2005). Our proposed framework falls in the first category and hence we will review some of relevant methods in the literature. Some methods use a simple position weight matrix (PWM) with a cut-off threshold and a similarity metric, e.g., MatInspector (Quandt *et al.*, 1995) and MATCH (Kel *et al.*, 2003), while others propose complicated PWMs in which the dependency among nucleotide positions is taken into account, e.g., di-nucleotide PWM (Siddharthan, 2010) and tree-based PWM (Bi *et al.*, 2011). Since these methods do not use prior knowledge on the location and the context of binding sites and merely depend on the DNA nucleotides, their prediction power is limited and this incurs quite low precision. The phylogenetic footprinting of TFBSs combined with PWM similarity metrics have been used to improve the prediction performance of TFBSs that are assumed to be conserved across orthologous species (Broos *et al.*, 2011; Sebestyén *et al.*, 2009; Tokovenko *et al.*, 2009; Marinescu *et al.*, 2005; Moses *et al.*, 2004). However, these methods still require a cut-off threshold to decide on putative binding sites. We recently proposed a threshold-free system, named FSCAN (Wang *et al.*, 2016), that utilizes the evolutionary conservation information of DNA sequences in addition to MISCORE-based similarity metrics (Wang and Tapan, 2012) in a fuzzy logic framework in order to predict the exact location of TFBSs. Characteristics of the DNA 3D structure were used as a replacement for PWMs to represent the TF-DNA binding specificity and add meaningful insights on the context of protein-DNA interaction (Gunewardena *et al.*, 2006; Burden and Weng, 2005; Oshchepkov *et al.*, 2004; Liu *et al.*, 2001; Ponomarenko *et al.*, 1999; Karas *et al.*, 1996). Features were extracted from DNA structural profiles and PWM similarity scores were used to build classifier models that recognize TFBSs efficiently and effectively. Perceptron neural networks (Steffen *et al.*, 2002), support vector machines(Bauer *et al.*, 2010), Bayesian networks (Nikolajewa *et al.*, 2007; Pudimat *et al.*, 2005), conditional random fields(Meysman *et al.*, 2011) and random forests (Broos *et al.*, 2013; Hooghe *et al.*, 2012) are among the machine learning algorithms that were used to build these classifiers.

In this paper, a threshold-free and machine learning-based framework, denoted by DNNESCAN, is proposed to recognize TFBSs in intergenic DNA sequences. The most recent ensemble learning technique DNNE (Alhamdoosh and Wang, 2014) is used to create classifier models for 22 TF proteins from the Saccharomyces Cerevisiae genome. The training and testing datasets are systematically extracted from genome-wide ChlP-chip experiments (Harbison *et al.*, 2004). Binding sites are characterized with base readout as well as shape readout features that help maximize the classification boundary between TFBSs and background sequences. DNNESCAN employs an efficient filtering procedure and oversampling algorithm to reduce the amount of background sequences and mitigate the high data imbalance ratio in training datasets, respectively. A filter-based features selection algorithm is also utilized to help alleviate the complexity of classifier models and hence improve the prediction performance. Finally, DNNESCAN predicts TFBSs at one nucleotide-level accuracy and embeds phylogenetic conservation knowledge in K-mer representation.

## 2 MATERIALS AND METHODS

### 2.1 Formulation and Notation

Given a set of intergenic DNA sequences 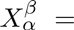 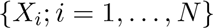 that are bound by a TF protein *β* under specific growth condition *α* in a ChlP-chip experiment. Our objective is to find the locations of binding sites in 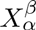 and other genomic regions that could attract the TF protein *β*. It is assumed that a prior knowledge on the DNA binding specificity of *β* is given in the form of a PWM and some binding sites in 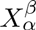 are already located. The forward and reverse strands of each input sequence should be searched for putative TFBSs. Each strand sequence of length *L* is partitioned into *L* − *K* + 1 K-mers *X_ij_* and only potential binding sites are reported. *X_ij_* is a short DNA sequence of length *K* (*K* is the width of PWM) that starts at position *j* and in the direction 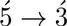. A particular DNA nucleotide at position *p* in a sequence *X_i_* is represented by 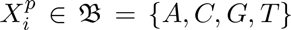 and the TF motif matrix PWM is denoted by *M_w_*. The rows of the matrix represent the four possible DNA nucleotides and the columns represent the binding site nucleotide positions that are zero-based indices. It is worth mentioning that PWM entries encode the log-odds probabilities of DNA base frequencies at each position in the TF motif along with genome background frequencies. However, PWM is sometimes unavailable and a position frequency matrix (PFM) is given instead. PFM holds only the frequencies of DNA nucleotides and can be easily converted to PWM for a given genome. Following (Wang and Tapan, 2012), a K-mer *X_ij_* is described using a 4 × *K* binary matrix *k* defined as follows

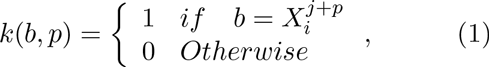

where 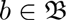 and *p* ∈ [0, *K*].

### 2.2 Learner models

Ensemble learning techniques have motivated many researchers to develop computational models in bioinformatics (Hooghe *et al.*, 2012; Bauer *et al.*, 2010). For example, the PhysBinder tool uses an ensemble learning technique called random forests to build TFBS classifiers (Broos *et al.*, 2013). We have recently proposed a new ensemble learning algorithm named DNNE (decorrelated neural networks ensemble) that outperforms the state-of-the-art ensemble learning algorithms in several regression problems (Alhamdoosh and Wang, 2014). Basically, an ensemble model is made up of *M* component base models and these base models are combined so that collective predictions are obtained. The advantage of ensemble learning over traditional learning algorithms such as neural networks and decision trees is that ensemble models exploit more regions in the feature space and thus have better generalization capabilities than single models (Hansen and Salamon, 1990).

In this paper, DNNE is used to learn classification boundaries for TFBSs. DNNE is a neural networks ensemble in which base models are random vector function link (RVFL) networks (Alhamdoosh and Wang, 2014) and they all have the same number of hidden neurons *L*. An RVFL network is a single layer feed-forward network whose input weights (between the input layer and the hidden layer) and hidden layer biases are assigned randomly and independently from the training data (Igelnik and Pao, 1995). The output weights (between the hidden layer and the output layer), however, are estimated based on the available training examples. The activation functions (called basis functions) of hidden neurons can be any squashing function and the output layer activation functions are linear. DNNE implements a novel analytical solution in order to calculate the output weights of ensemble base networks. The solution is based on normal equations and singular value decomposition techniques. An estimation for the global output matrix 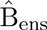 is calculated using the following formula

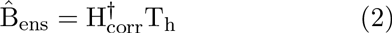

where H_corr_ is *ML*×*ML* hidden correlation matrix, 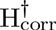 is the generalized pseudo-inverse of H_corr_, T_h_ is *ML* × *n* hidden-target matrix, and *n* is the number of ensemble model outputs. Calculating H_corr_ and T_h_ requires a regularizing factor λ ∈ [0,1]. More details on how to calculate them can be found in (Alhamdoosh and Wang, 2014). As DNNE is used in this paper for a classification task, the majority voting technique is employed to make classification decisions. In other words, the predicted class label (binding site +1 or background K-mer −1) is assigned as the most commonly predicted class label by base RVFL networks. The classification decision of the DNNE model is given by the following formula

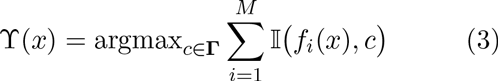

where *f_i_*(*x*) is the output of the ith RVFLbase network when an instance *x* is presented, **Γ** is the set of class labels and 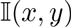 is the identity function, i.e., 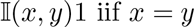.

Three parameters need to be set in order to optimize a DNNE model, i.e., the base model size *L*, the ensemble size *M* and the regularizing factor λ. From our observations in (Alhamdoosh and Wang, 2014), *M* is sufficient to be between 3 and 9 and it is recommended to select an odd number, λ is recommended to be 0.55, and *L* plays a key role in the learning and generalization capabilities of the DNNE model. Usually, random weight networks, like DNNE models, require a large number of hidden neurons *L* though some data can be successfully modelled using a few hidden neurons only (Alhamdoosh and Wang, 2014).

### 2.3 Biological data

In order to assess the performance of our proposed framework, we collected ChIP-chip sequence-sets for 203 verified TFs from the budding yeast genome. (Harbison *et al.*, 2004) investigated the binding preferences of each TF protein over around 6,000 intergenic regions (IGRs), which cover the whole genome, and under 14 different growth conditions. As a result, 350 experiments were conducted and each one produced a sequence set of the form 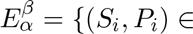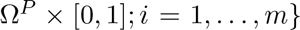, where *P_i_* is p-value (the binding probability of a protein *β* to a probe sequence *S_i_* under growth condition *α*) and Ω*^P^* is a set of *m* probes. (MacIsaac *et al.*, 2006) reanalyzed these sequence-sets and discovered motif matrices (PWM) and putative binding sites for 124 TF proteins. The Saccharomyces Genome Database (SGD) adopts the TFBSs produced by a later study as one its transcription regulation tracks. In this paper, we update the start and end positions of probe and IGR sequences according to the genome release *R*64.1.1, published in February 2011 at SGD (Cherry *et al.*, 2012) and (MacIsaac *et al.*, 2006)’s TFBSs for the same release are also obtained. Furthermore, we consider the whole IGRs overlapping with bound probes and of length more than 30bp in order to discover more potential TFBSs. The mapping of probes on IGRs along with their potential regulated target genes were obtained from (Harbison *et al.*, 2004)’s supplementary data.

#### 2.3.1 Preparing sequence-sets

Two sequence-sets are required for each ChIP-chip experiment 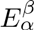: bound sequences and unbound sequences. The bound sequence-set is defined as 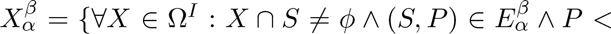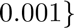 where Ω*^I^* denotes the set of all IGRs in the genome, and the unbound sequence-set is given by 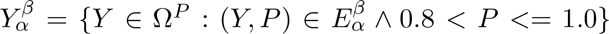. The cardinality of 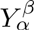 should be at least five times of the cardinality of 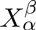 so that the probes of largest p-value are taken first. Now, we define selection criteria for the bound sequence-sets used in this study as follows

- *β* is a TF with a known binding motif matrix (PFM),
- 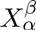 contains only IGRs that regulate verified open reading frames,
- 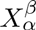 has at least 50 known binding sites in SGD,
- And the cardinality of 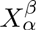 is greater than or equal to 50.

Only 38 out of the 350 sequence-sets satisfy the above criteria and they correspond to 22 TF proteins. The sequence-set with the largest number of *known* binding sites is selected for each TF. Table 1 shows the number of IGR sequences, average sequence length, total number of nucleotides (one strand), number of known TFBSs and the width of PWM for the selected 22 sequence-sets. The sequence-set name is denoted by the TF gene name and the experiment growth condition. PWMs are downloaded from (MacIsaac *et al.*, 2006)’s supplementary materials and converted into PFMs as needed by DNNESCAN in three steps: (i) the frequencies *P_b_* of *A, C, G, T* in Ω*^I^* are calculated, (ii) PFM entry of base *b* and position *p* is calculated *P_f_*(*b,p*) = *P_b_* × 2*^P_w_(b,p)^*, and (iii) PFM columns are normalized by dividing its elements by Σ_*b*_ *P_f_* (*b, p*). Finally, the motif matrices are trimmed based on the IUPAC consensus sequences (Cavener, 1987) of the corresponding TF. The head and tail positions that correspond to ‘.’ (any) in the consensus sequences are trimmed.

**Table 1:**
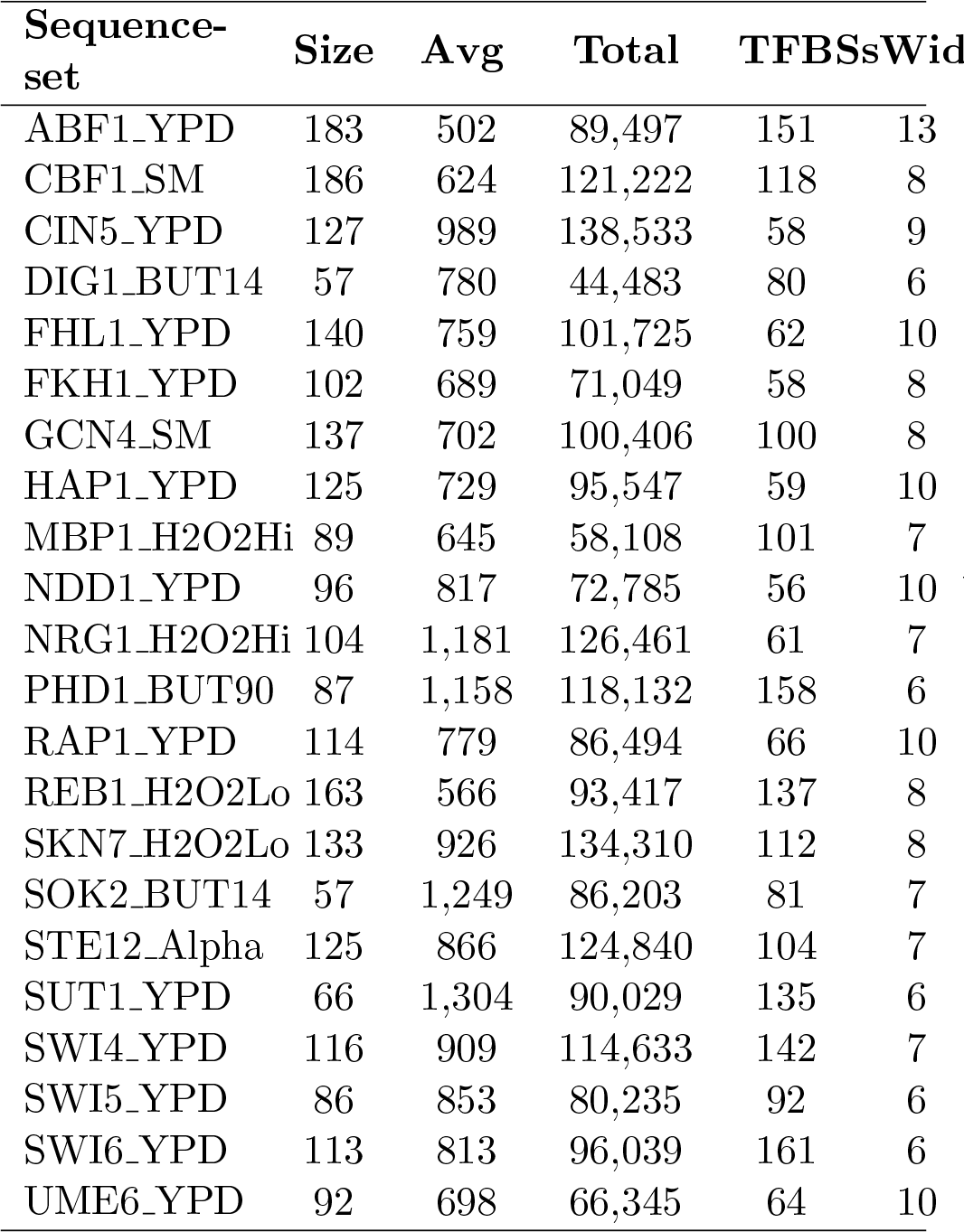
*Details of the bound sequence-sets 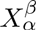 for each investigated TF.*

### 2.4 Features extraction

In a machine learning based framework, objects need to be described by some numerical or nominal features. K-mers and their anking regions are the main objects for any TFBS predictor. In this paper, we characterize true binding sites as well as background K-mers with many numerical features that can be categorized into four groups based on their nature and function. DNNESCAN utilizes base read-out, shape readout and evolutionary characteristics of DNA sequences in order to model TF protein-DNA binding specificities. Next, we explain how to calculate each one of these features.

#### 2.4.1 Motif-dependent features

**Mean and width of binding affinity:** The strength of DNA-protein binding strongly depends on which amino acids in the protein contact which DNA nucleotides in the DNA sequence (Luscombe and Thornton, 2002). However, some amino acids in the DNA-binding domain of TF proteins strongly bind to the binding site bases while others weakly bind to their corresponding bases (Zhao *et al.*, 2012). This motivated us to define a similarity metric between a K-mer and a PFM so that not all positions in the PFM and the K-mer sequence are included 1ththe similarity score. *R* positions are randomly selected from PFM with their corresponding bases in the K-mer. Then, inspired by (Wang and Tapan, 2012), the similarity between a K-mer matrix *k* and a PFM *M_f_* is calculated only using the randomly selected positions *R_p_* as follows

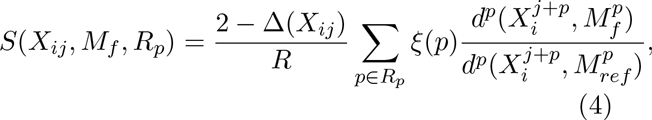

where 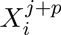, 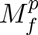, and 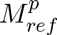 are the *p*th positional columns in *X_ij_*, *M_f_*, and *M_ref_*, respectively, *M_ref_* is the PFM of the background reference model and *d^p^*() is a special generalized Hamming distance function that measures the dissimilarity between a PFM and a K-mer at a specific position, as follows

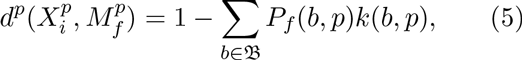

*ξ*(*p*) is the degree of conservation of position *p* in the *M_f_* matrix and is given by the information entropy

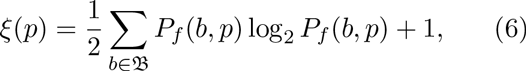

and Δ(*X_ij_*) represents the over-representation of K-mer *X_ij_* in the bound sequence-set *X* and is defined by the following formula

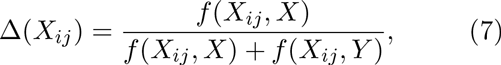

where *f*(*z*, *Z*) is the frequency of K-mer *z* in the sequence-set *Z*.

The random sampling of positions is repeated T times and a *DNA binding affinity signal* (DNA-BAS) is generated for each K-mer accordingly. K-mers that could be putative binding sites would have quite low amplitude DNA-BAS while non-functional K-mers would have very high amplitude (Wang *et al.*, 2016). To characterize DNA-BAS, the *mean* is defined as

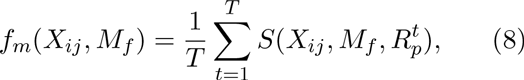

where 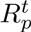 is the set of randomly selected positions at trial *t*, and the *width* is defined as

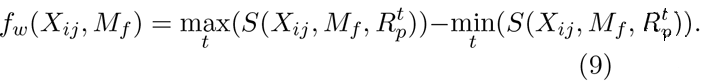

The number of selected positions *R* is usually set to 60% of the motif width, that is, *R* = [0.6 × *K*] and the number of random trials *T* is usually set to 100 (Wang *et al.*, 2016).

**PWM constraint**: This feature directly measures the degree of conservation of a K-mer with respect to a PWM (Fu *et al.*, 2009) and is given by

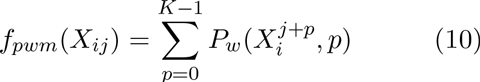

where *P_w_*(*b,p*) is the log-odds value of the base *b* at position *p*. The discriminative power of this feature mainly depends on the quality of the supplied PWM/PFM.

**Conservation symmetry:** This feature is constructed based on the empirical observation that the DNA binding domain of TF proteins have symmetric conservation profiles around the centers of their binding sites (Fu *et al.*, 2009). This symmetry is captured in a given K-mer as follows

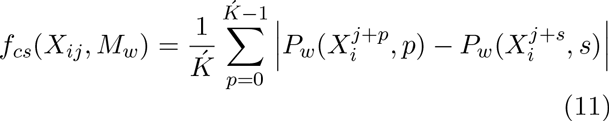

where *s* = *K* ‒ *p* ‒ 1 and 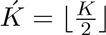.

#### 2.4.2 Structural profiles features

**Conformational and thermodynamic features:**Though the chemical mosaic of DNA bases of binding sites plays an important role in TF protein-DNA binding, the 3D structure of a putative TFBS and its flanking sequences control the quality of binding specificity for many TF proteins. These shape readout interactions between protein and DNA macromolecules particularly help distinguish between the specificities of TF proteins that belong to the same structural family (Rohs *et al.*, 2010). Conformational and physiochemical aspects of DNA structure are encoded in dinucleotide or trinucleotide profiles that could be used for sequence analysis applications (Hooghe *et al.*, 2012; Meysman *et al.*, 2011; Baldi and Baisnee, 2000). Dinucleotide properties from the DiProDB database are used in this paper (Friedel *et al.*, 2009). DiProDB has more than 100 dinucleotide structural profiles of DNA sequences and each is represented by 16 numerical values that ccifrespond to all possible dinucleotide conformations. Sihtre. most of these profiles are quite correlated, we propose a simple algorithm to select dinucleotide properties that convey more than 99% of the information in DirProDB and well characterize the 3D structure of a given DNA sequence. Initially, 103 conformotional and physiochemical dinucleotide profiles were obtained from DiProDB and grouped based on their names. The profile of the most recently published was selected from each group and hence 65 profiles are collected. 49 profiles describe conformational properties of DNA and 16 describe physiochemical properties. Finally, our general structural models selection (GSMS) algorithm is applied on each property category independently. 9 out of the 49 conformational profiles are selected: major groove distance, clash strength, tilt_rise, rise, tip, twist_shift, twist_rise, direction, and inclination, and 6 out of the 16 thermodynamic profiles are selected: flexibility_slide, melting temperature, slide stiffness, flexibility_shift, stacking energy, mobility to bend towards major groove. Noticeably, most of the selected structural models have been used in TFBS prediction frameworks, i.e., PhysBinder (Broos *et al.*, 2013) and CRoSSeD (Meysman *et al.*, 2011). Algorithm 1 describes GSMS procedure. The algorithm requires the structural profiles to be organized in a 4*^m^* × *S* matrix 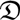 where *S* is the number of profiles and *m* is the order of profile oligonucleotides. GSMS uses principal component analysis (PCA) (Pearson, 1901) to select structural profiles that cover some percentage of the total variance in the whole DiProDB. The correlation matrix is used here because dinucleotide structural properties have different scales.

Now, we can characterize TFBSs with numerical features based on the above selected profiles. Since the structural characteristics of a DNA macromolecular cannot be perceived in short helices (¡ 45 bp) like TFBSs (Rohs *et al.*, 2010), the left and right flanking regions of a given K-mer are considered in the feature extraction. The average score of a given dinucleotide property 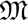 for a given K-mer *X_ij_* is calculated as follows

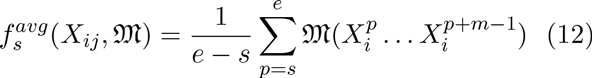

where *s* = *j ‒ w, e = j* + *K* + *w ‒ m* + 1, *e* > *s, w* is the size of the left and right flanking regions, and 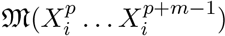 is the measurement of profile 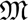 for the oligonucleotide 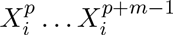. Flanking regions of size *w* = 20*bp* are used in DNNESCAN. Furthermore, oligo-based structural features are calculated for a DNA sequence in order to measure the contribution of oligonucleotides in each aspect of the DNA 3D structure. These features consider the over-representation of oligonucleotides in the K-mer and its flanking regions. Since structural profiles are estimated from double-stranded DNA sequences, 10 (instead of 16) unique oligo-based features are calculated for each dinucleotide property as follows

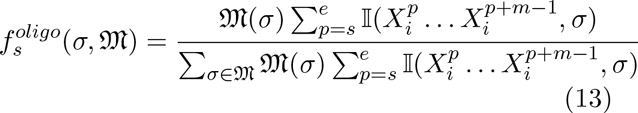

where *σ* is an oligonucleotide and 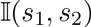 is the identity function. At the end, each K-mer is encoded with 165 conformational and physiochemical features.

**Algorithm 1.**
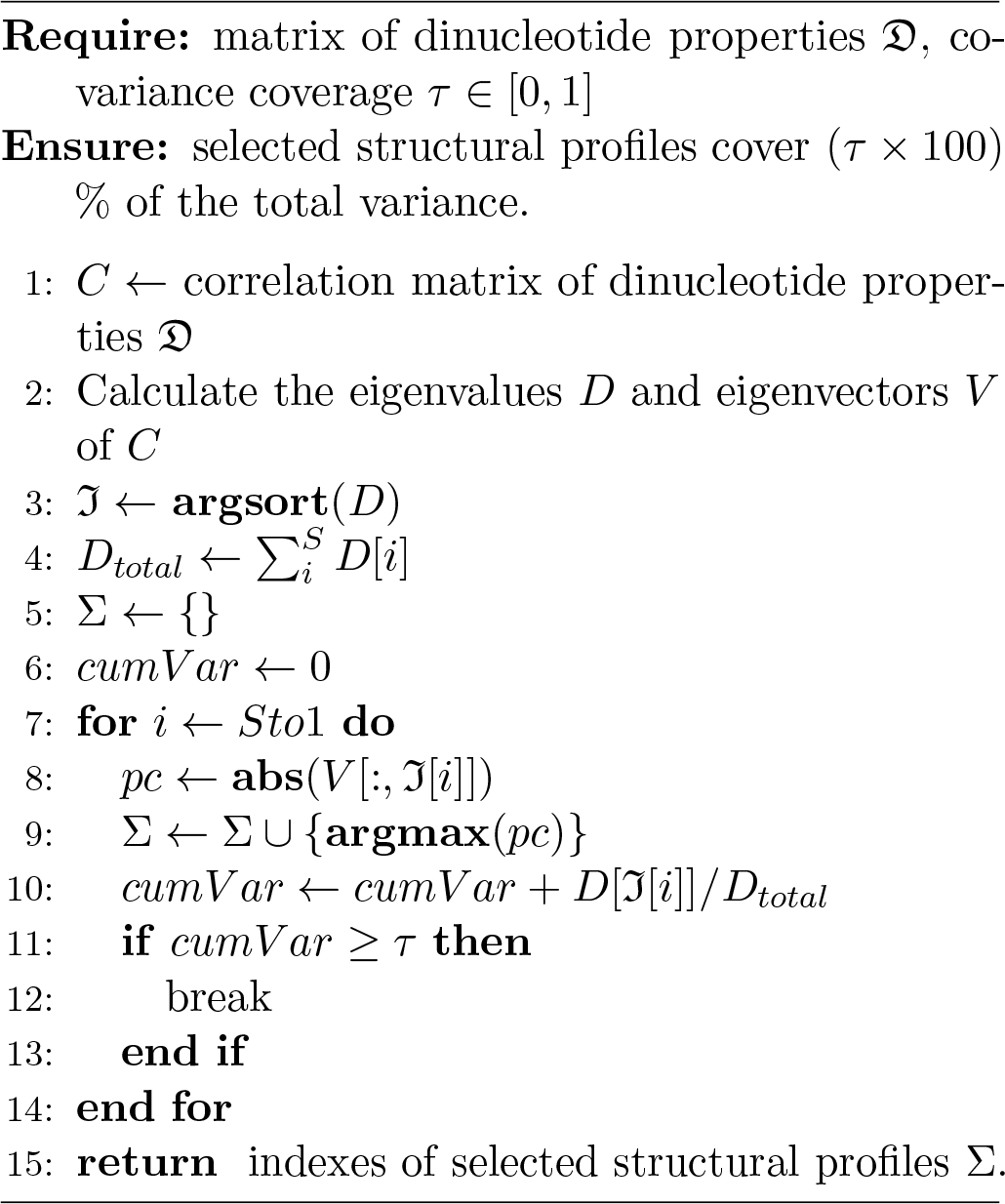
General structural models selection algorithm (GSMS).

**Simple nucleosome occupancy:** The positioning of nucleosomes along the DNA is believed to provide a mechanism for differential access to TFs at potential binding sites. It has been shown that the functional binding sites of TFs at regulatory regions are typically depleted of nucleosome (Narlikar *et al.*, 2007). We used the computational model published by (Kaplan *et al.*, 2009) to predict the probability of each nucleotide position in the yeast genome being bound by a nucleosome. Then, the nucleosome occupancy score for a K-mer *X_ij_* is calculated as follows

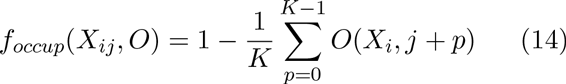

where *O*(*X_i,j_*) is the probability of the *j*th position in sequence *X_i_* being occupied by a nucleosome.

#### 2.4.3 Letter-based features

**Sequence composition features:** These features reflect the nucleotide composition of a sequence. To achieve this, we apply the GSMS algorithm on the letter-based dinucleotide properties in DiProDB (Friedel *et al.*, 2009). We ended up with 4 letter-based profiles that cover more than 99% of the total variance in this group. They are: purine content, GC content, Guanine content and Keto content. The prevalence of Guanine and Cytosine in a genomic regions indicates that they may contain regulatory elements such as TFBSs since DNA structures with high GC-content tend to have higher stability (Fu *et al.*, 2009). Equation 12 is used to extract K-mer features using the selected letter-based dinucleotide profiles.

**Reverse complementarity:** This feature measures the similarity of a potential binding site *X_ij_* to its counterpart on the other genomic strand (Fu *et al.*, 2009). It measures the similarity around the center of a motif model as follows

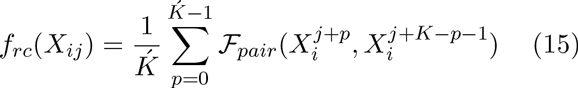

where 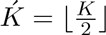 and *Ϝ_pair_*(*a, b*) produces 1 iff *a* and *b* are a Watson-Crick pair.

**CpG Island Occurrence:** This feature determines the existence of CpG-rich regions in a K-mer and its neighborhood using the observed-to-expected CpG ratio and the GC-content(Gardiner-Garden and Frommer, 1987). It measures the probability of K-mer occurrence in a CpG island. Observed-to-expected CpG ratio is calculated as follows

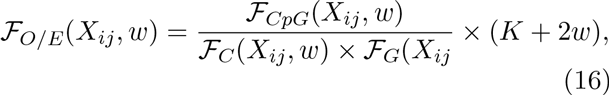

where *w* is the flank size, *Ϝ_CpG_*(*X_ij_,w*) gives the number of CpG dinucleotides in *X_ij_* and its flanks, and *Ϝ_C_*(*X_ij_,w*) and *Ϝ_G_*(*X_ij_,w*) give the number of Cytosines and Guanines in *X_ij_* and its flanks, respectively. Therefore, the CpG island occurrence feature is defined as

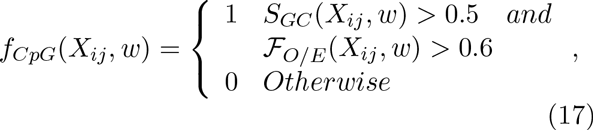

where *S_GC_*(*X_ij_,w*) is the GC content ratio in *X_ij_* and its flanks.

#### 2.4.4 Phylogenetic footprinting feature

The rationale behind this feature is that genomic regions that contain TFBSs have very strong phylogenetic relationships (Blanchette and Tompa, 2002). For this purpose, the conservation scores for the S. Cerevisiae are obtained from the UCSC genome browser. These scores are produced for each base in the genome using the phastCons program that utilizes the multiple sequence alignment of 6 yeast genomes with the S. Cerevisiae genome: *S. Cerevisiae, S. Paradoxus, S. Mikatae, S. Kudriavzevii, S. Bayanus, S. Castelli* and *S. Kluyveri.* To convert these conservation scores into a discriminative feature that characterizes conserved TFBSs, a simple function is defined as follows

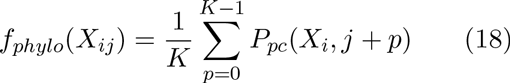

where *P_pc_*(*X_i_,p*) is the phastCons score at position *p* in sequence *X_i_*. This feature plays a key role in discriminating functional binding sites from non-functional K-mers that have the same DNA nucleotide sequence but occur in different genomic regions.

### 2.5 Filtering and classifiers

The current learning-based approaches of TFBS prediction use two types of data to build classification models: a positive example set solely based on the known binding sites and a negative example set randomly created from different regions of the genome.

These approaches use a high cut-off threshold to filter out background K-mers (Hooghe *et al.*, 2012; Meysman *et al.*, 2011). The main reason behind this procedure is to reduce the imbalance ratio between positive and negative examples that could reach as high as 1:5000 in a ChlP-chip sequence-set (Wang *et al.*, 2016). Since threshold-based filtering is experimental and has no solid foundation, DNNES-CAN uses robust filtering algorithms that maintain background filtering, data imbalance reduction, and feature selection as discussed in the next sub-section.

#### 2.5.1 Two-stage background filtering

The excessive redundancy of background DNA sequences makes building classification models for TFBS prediction very challenging. Therefore, removing unnecessary DNA sequences is highly desirable and would facilitate the learning of decision boundaries. Similar to FSCAN (Wang *et al.*, 2016), DNNESCAN uses a two-stage filtering procedure. In the first stage, either the forward or reverse strand of a K-mer is retained depending on the smaller value of the mean of DNA-BAS *f_m_*. In the second stage, however, an efficient filter based on the mean and width of DNA-BAS is developed to further remove background K-mers. Our proposed algorithm, called adaptive ellipsoidal filter (AEF), learns an ellipsoidal boundary for the true binding sites in the 2D space (*f_m_, f_w_*) ∈ ℝ^2^ and then uses this boundary to filter non-functional K-mers. The AEF algorithm greatly reduces the *BG/BS* imbalance ratio with only as low as 1% of the whole true binding sites are wrongly filtered out (Wang *et al.*, 2016). The novelty of AEF emerges in its ability to build a customized filter for each TF protein rather than using one cut-off threshold for all TF proteins.

#### 2.5.2 Imbalance reduction using RANDOVER

After applying AEF on a ChIP-chip sequence-set of a given TF protein, the resulting K-mers set is still highly biased towards the background sequences. This causes any traditional learning algorithm (e.g., DNNE) to fail to learn the underlying data distribution and therefore generate classification models with very large learning and generalization errors. In order to overcome this limitation and obtain a well-balanced training dataset, an algorithm called RANDOVER (random oversampling) is proposed earlier to control the imbalance ratio of a K-mer set (Wang *et al.*, 2016). RANDOVER is a simple and efficient solution that adds artificial training examples labeled with a minority class label. The newly added examples are sampled from the existing minority examples and then perturbed with a small amount of noise as described in (Wang *et al.*, 2016).

#### 2.5.3 Features selection

The dimensionality of the K-mer features vector is another factor that affects the learning capability of a classifier model and hence reduces its predictive power as dimensionality increases (Saeys *et al.*, 2007). In DNNESCAN, K-mers are distributed in a 270-dimensional space though only few of these dimensions are sufficient to model the protein-DNA binding specificity for most TFs. Feature selection algorithms were proposed to find an optimal subset of features that is least redundant and most informative. Therefore, the underlying class distribution and the classifier models would be quite simple and easy to interpret and understand. Wrapper-based feature selection was used recently in PhysBinder to build TFBS random forest classifiers (Broos *et al.*, 2012). On the other hand, filter-based methods can be used as an alternative to wrappers since their computational complexity is much lower (Blum and Langle, 1997). Filter techniques basically rank features according to their relevance to the class labels and remove irrelevant and redundant features accordingly. DNNESCAN employs an efficient and effective filter algorithm called Fast correlation based filter (FCBF) (Yu and Huan, 2003). FCBF uses the concept of predominant feature to build the optimal subset of features. Predominant feature is a feature that has information about the class distribution and also about all other features that are Markov blanket by this predominant feature. It takes into account the relevance and redundancy of features so that the selected features have very high class-feature correlation and low feature-feature correlation (Yu and Liu, 2004). FCBF uses the symmetrical uncertainty (SU) to measure the correlation between features (redundancy) and between features and class (relevance). Obviously, the *SU* measure requires nominal or discrete feature values rather than continuous values. Since all the K-mer features in DNNESCAN are continuous, an efficient discretization algorithm that uses an information entropy minimization heuristic is applied to convert continuous features into multiple intervals features (Liu *et al.*, 2002). Linear correlation measures, e.g., Pearson correlation, cannot capture nonlinear relationships between features. Therefore, information theory concepts, e.g., entropy, are used to measure these correlations. (Fayyad and Irani, 1993) proposed splitting a continuous range of feature-values into multiple intervals using the minimum description length principle (MDLP). The algorithm uses MDLP to decide on the partitioning of intervals. To discretize a continuous attribute, the proposed algorithm first sorts the attribute values in an ascending order. Then, it recursively splits the attribute range of values into two intervals based on the best class entropy of a cut-point. The recursive splitting is terminated when (Fayyad and Irani, 1993)’s MDLP criterion is met.

### 2.6 DNNESCAN: a framework for TFBS Prediction

DNNESCAN framework is composed of two main parts. The first part (*building module*) is used to build TFBS classification models using a given training sequence-set and the second part (*testing module*) is to identify TFBSs using the learned classifiers by scanning DNA sequences. The building module is made up of a pre-processing unit and learning unit while the testing module has only the classification unit that implicitly utilizes some of the functions of the pre-processing and learning units. The first step in the pre-processing unit is K-mer extraction. A window of size *K* is used to scan DNA sequences and generate K-mers by shifting the window one nucleotide position each time. As a result, a sequence of length *L* is partitioned into *L* ‒ *K* +1 K-mers. The reverse complement of each K-mer is generated as well. Next, the width and mean of a DNA binding affinity signal are calculated for each K-mer and its reverse complement using the PFM model. The two-stage background filtering is applied to reduce the number of non-functional K-mers in the sequence-set. In the training module, an adaptive ellipsoidal filter model is built using the training K-mers and then it is used to filter out background K-mers in the training and testing sequences. Once the unnecessary K-mers are filtered out, a features extraction procedure is executed to calculate the numerical features for each of the remaining K-mers as was thoroughly explained earlier. The values of these features are then normalized to [0, 1] to remove the bias of different feature scales.

Once K-mers are characterized with numerical features, the DNNE models can be learnt from these labeled feature vectors. However, the data imbalance reduction algorithm is needed to reduce the imbalance between true binding sites and background K-mers. Afterward, the feature selection algorithm is applied and the selected features are decided based on the training K-mer set only to be used for testing K-mers as well. Eventually, neural network ensemble models are created using the DNNE learning algorithm (Alhamdoosh and Wang, 2014) and a K-mer is classified as a true binding site if the DNNE output is greater than 0 and as a background sequence otherwise. Moreover, all K-mers that are filtered out in the pre-processing stage are labeled as background sequences. Fig. 1 illustrates the architecture of our proposed framework.

**Figure 1:**
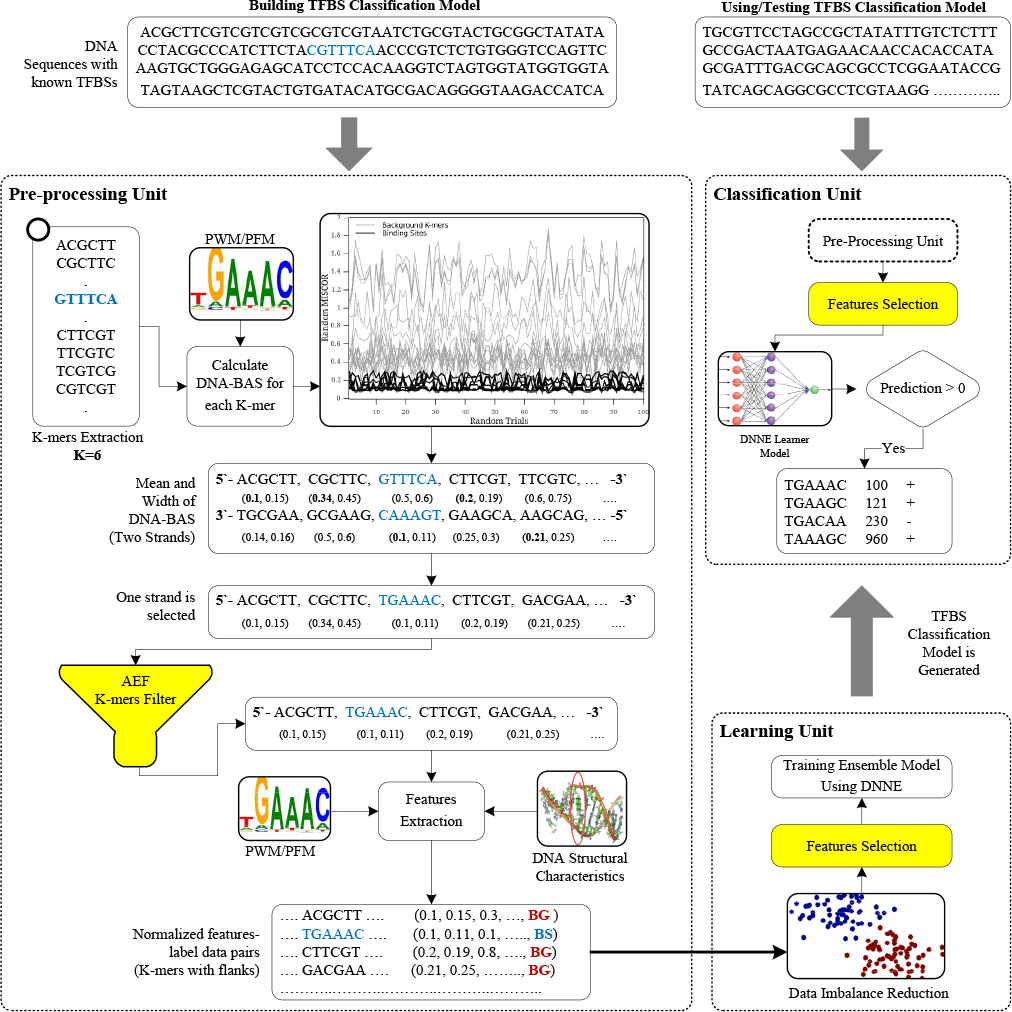
*Components and functionalities of DNNESCAN system.*

### 2.7 Performance evaluation

The 10-fold cross-validation procedure is conducted to evaluate the performance of DNNESCAN and FSCAN on each ChlP-chip sequence-set. Each sequence-set is divided into 10 subsets so that each subset contains an equal number of background K-mers and true binding sites. The 10 subsets initiate 10 runs for each sequence-set in which each subset is tested once and used more than once for training or validation. For each run, two subsets are selected for testing and validation and the remaining subsets are combined together to form the training sequence-set. A 2 × 2 confusion matrix is created based on the labeled testing K-mers and 10 confusion matrices produced from the 10 runs are summed in order to form one final confusion matrix. Each experiment is repeated 10 times to alleviate the bias of crossvalidation partitioning and DNNE random weights. On the other hand, the performance of MatInspector (Quandt *et al.*, 1995) is assessed on all possible cut-off thresholds in [0.7,1.0] and the best performance measured using the F1-measure is reported. Actually, different metrics can be used to evaluate the performance of TFBSs predictors (Tompa *et al.*, 2005). We use four performance indexes that are commonly used in machine learning practice to evaluate classification systems: precision (P) to measure the exactness, Recall (R) to measure the completeness and F1-measure to obtain a meaningful insight on precision and recall together. It is worth mentioning that we count a K-mer correctly predicted as a binding site if its location exactly matches a known site location regardless of the strand. Since the performance of DNNESCAN is evaluated on a collection of sequence-sets, a system success ratio (SSR) is defined to compare the performance of two systems *A* and *B* as follows

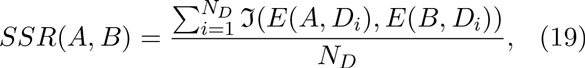

where *E*(*A,D_i_*) is the evaluation metric of system *A* on a sequence-set *D_i_*, 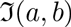 returns 1 iff *a >= b,* and *N_D_* is the total number of sequence-sets.

## 3 RESULTS and DISCUSSION

### 3.1 Performance Analysis of DNNESCAN

In this section, we thoroughly analyze the performance of DNNESCAN using the 22 ChIP-chip sequence-sets. Table 2 shows the average precision, recall and F1-measure with the standard deviation for DNNESCAN on each of these datasets. It also shows the best performance for DNNESCAN over multiple experiment repetitions. It is clear that our DNNESCAN has relatively high recall for most of the datasets. It shows a recall higher than 75% on 14 out of the 22 datasets while it shows a recall between 60% and 75% on the other 8 datasets. The overall average recall over all datasets is 77% while the average best recall is 79%. Furthermore, the sensitivity of our system against false positives seems to be good. DNNESCAN demonstrates a precision higher than 60% on 4 datasets (ABF1, REB1, SWI4 and UME6) and less than 50% on 9 datasets while the system precision ranges between 50% and 60% on the other 9 datasets. This low positive predictive value of DNNESCAN could be ascribed to two factors. First, a lack of known TFBSs in the tested DNA sequences causes true TFBSs to be predicted as false positives. We show in Section 3.3 that some of the binding sites discovered by DNNESCAN have a very high potential to be true regulatory sites. Second, the quality of data that has been used for building DNNE models plays a key role as well. DNNESCAN shows 49% average precision over all sequence-sets with 51% in the best situation.

**Table 2:** *Performance summary of DNNESCAN on 22 ChIP-chip sequence-sets.*

| Dataset | TFBS Prediction |  |  |  |  |  | Number of<br>Selected<br>Features |
| --- | --- | --- | --- | --- | --- | --- | --- |
|  | Precision |  | Recall |  | F1-Measure |  |  |
|  | Average | Best | Average | Best | Average | Best |  |
| ABF1_YPD | 0.74 ± 0.01 | 0.75 | 0.96 ± 0.01 | 0.97 | 0.84 ± 0.01 | 0.85 | 2.14 ± 0.02 |
| CBF1_SM | 0.58 ± 0.01 | 0.59 | 0.71 ± 0.02 | 0.73 | 0.64 ± 0.01 | 0.65 | 2.35 ± 0.02 |
| CIN5_YPD | 0.3 ± 0.02 | 0.34 | 0.75 ± 0.01 | 0.74 | 0.43 ± 0.02 | 0.47 | 2.03 ± 0.01 |
| DIG1_BUT14 | 0.52 ± 0.02 | 0.54 | 0.82 ± 0.02 | 0.84 | 0.64 ± 0.01 | 0.66 | 3.26 ± 0.03 |
| FHL1_YPD | 0.33 ± 0.03 | 0.36 | 0.66 ± 0.03 | 0.69 | 0.44 ± 0.03 | 0.48 | 2.49 ± 0.03 |
| FKH1_YPD | 0.55 ± 0.02 | 0.56 | 0.76 ± 0.03 | 0.81 | 0.63 ± 0.02 | 0.66 | 2.12 ± 0.02 |
| GCN4_SM | 0.57 ± 0.01 | 0.57 | 0.75 ± 0.02 | 0.79 | 0.65 ± 0.01 | 0.66 | 2.75 ± 0.02 |
| HAP1_YPD | 0.49 ± 0.01 | 0.51 | 0.92 ± 0.03 | 0.95 | 0.64 ± 0.02 | 0.66 | 2.21 ± 0.02 |
| MBP1_H2O2Hi | 0.48 ± 0.05 | 0.5 | 0.53 ± 0.05 | 0.61 | 0.5 ± 0.04 | 0.55 | 4.42 ± 0.06 |
| NDD1_YPD | 0.27 ± 0.02 | 0.28 | 0.78 ± 0.02 | 0.82 | 0.4 ± 0.02 | 0.42 | 3.42 ± 0.04 |
| NRG1_H2O2Hi | 0.56 ± 0.02 | 0.56 | 0.73 ± 0.03 | 0.79 | 0.63 ± 0.02 | 0.66 | 2.64 ± 0.04 |
| PHD1_BUT90 | 0.49 ± 0.01 | 0.5 | 0.72 ± 0.02 | 0.74 | 0.58 ± 0.01 | 0.6 | 2.03 ± 0.01 |
| RAP1_YPD | 0.42 ± 0.02 | 0.47 | 0.78 ± 0.03 | 0.82 | 0.55 ± 0.02 | 0.6 | 3.61 ± 0.05 |
| REB1_H2O2Lo | 0.82 ± 0.01 | 0.83 | 0.92 ± 0.01 | 0.93 | 0.87 ± 0.01 | 0.88 | 4.88 ± 0.08 |
| SKN7_H2O2Lo | 0.32 ± 0.02 | 0.35 | 0.88 ± 0.02 | 0.92 | 0.47 ± 0.02 | 0.51 | 2.49 ± 0.04 |
| SOK2_BUT14 | 0.3 ± 0.01 | 0.31 | 0.8 ± 0.03 | 0.83 | 0.44 ± 0.01 | 0.46 | 2.89 ± 0.06 |
| STE12_Alpha | 0.39 ± 0.02 | 0.41 | 0.71 ± 0.03 | 0.74 | 0.51 ± 0.02 | 0.53 | 3.44 ± 0.04 |
| SUT1_YPD | 0.36 ± 0.01 | 0.37 | 0.82 ± 0.01 | 0.82 | 0.5 ± 0.01 | 0.51 | 2.97 ± 0.02 |
| SWI4_YPD | 0.66 ± 0.01 | 0.68 | 0.75 ± 0.01 | 0.75 | 0.7 ± 0.01 | 0.71 | 4.64 ± 0.03 |
| SWI5_YPD | 0.36 ± 0.03 | 0.41 | 0.6 ± 0.03 | 0.62 | 0.45 ± 0.02 | 0.49 | 3.66 ± 0.06 |
| SWI6_YPD | 0.49 ± 0.01 | 0.51 | 0.61 ± 0.02 | 0.65 | 0.54 ± 0.02 | 0.57 | 4.62 ± 0.02 |
| UME6_YPD | 0.78 ± 0.01 | 0.79 | 0.88 ± 0.01 | 0.89 | 0.83 ± 0.01 | 0.84 | 2.19 ± 0.01 |
| Average | 0.49 ± 0.02 | 0.51 | 0.77 ± 0.02 | 0.79 | 0.59 ± 0.02 | 0.61 | 3.06 ± 0.03 |

It can be easily seen from Table 2 that DNNESCAN demonstrates an Fl-score higher than 60% for most of the testing datasets. Three datasets, namely ABF1, REB1 and UME6, confirm that DNNESCAN is quite accurate in recognizing TFBSs and its F1-measures on these datasets are 88%, 85%, and 84%, respectively. On the contrary, DNNESCAN performs badly on three datasets, namely CIN5, NDD1 and SOK2, with Fl-measures 42%, 46% and 47%, respectively. To understand under which conditions DNNESCAN performs effectively, we closely examine these six datasets. DNNESCAN shows relatively high sensitivity on these sequence-sets while its precision drops as low as 27% for some of them. On one hand, the ABF1 transcription factor has a long specificity motif with high binding specificity at its two ends that helps increase the positive predictive value of DNNESCAN. Further, DNEEScan shows the best F1-measure on the REB1 sequence-set whose TF is featured with quite a conserved binding motif and hence the width and mean of DNA-BAS would be quite small. As a result, the AEF background filter removes many false positives at the early stage of DNNESCAN system prediction. Similarly, the UME6 protein has rather high specificity to its binding site nucleotides and a relatively long motif. On the other hand, CIN5, DD1 and SOK2 have quite weak DNA motifs and low DNA binding specificity. This results in high binding affinity signals and produces abundant false positives that degrade DNNES-CAN precision. Table 3 shows that DNNESCAN binding models of CIN5, DD1 and SOK2 mainly depend on the PWM score and phylogenetic conservation features.

Next, we closely examine the K-mer features that were selected by the FCBF algorithm. The last column of Table 2 shows that the average number of selected features over 1000 experiments ranges between 2 and 4 for most datasets and only four datasets (MBP1, REB1, SWI4 and SWI6) require more than four features. In order to understand which features are selected for each TF protein, we list the top six selected features for each dataset along with their selection rates between [] in Table 3. The feature selection rate is the proportion of times a feature is selected per 1000 experiments and is computed as a percentage. It is apparent that only few features are necessary to characterize the DNA-binding affinity for most of the investigated TF proteins. The phylogenetic conservation and the PWM constraint features have been dominantly selected by DNNESCAN for seven TFs (see CBF1, CIN5, HAP1, NRG1, PHD1, SKN7 and UME6 in Table 3) and they demonstrate more than 97% selection rates. The other four top selected features appear in less than 20% of the simulations and mainly include structural features of K-mers and their flanking regions. It can be also noted from Table 3 that only specific di-nucleotides of the conformational and physiochemical profiles contribute to the DNA-binding affinity of these seven TFs. The DNA-binding specificity of ABF1 and FHL1 is dominated by two features only. The PWM constraint is always selected for both TFs while the twist_shift of the di-nucleotide CC/GG and the tip of AC/TG are highly preferred by ABF1 and FHL1, respectively. Furthermore, the guanine content and the twish_shift of AA/TT might play a significant role in the binding of the FHL1 protein to regulatory sites. Besides the phylogenetic conservation and the PWM constraint features, five TF proteins show great tendency towards DNA structure characteristics in their DNA-binding specificities. The DNA-binding affinity of DIG1, FKH1, GCN4, NDD1 and SUT1 seem to be leveraged by the slide stiffness of TA/AT, the mean of DNA-BAS, guanine content, purine content and clash strength of CA/GT, respectively. By examining the selection rates of DIG1, FKH1 and NDD1 features in Table 3, the DNA-binding affinity of these TFs could be attributed to the purine content in the putative binding region, the width of DNA-BAS and the rise of base pairs CC/GG, respectively. It can be also observed that several conformational and physiochemical features besides the phylogenetic conservation and PWM constraint significantly control the DNA-binding affinity of RAP1, SOK2, STE12, SWI4, SWI5 and SWI6. Finally, our results show two interesting cases. It can be seen from Table 3 that there is no prevailing feature characterizing the DNA-binding specificity of MBP1 and all the top six features seem to be important for MBP1 to bind on DNA regulatory elements. Similarly, DNNESCAN utilizes two motif-dependent features, three conformational features, and one letter based feature to model the DNA-binding affinity of REB1.

**Table 3:** *The top six features selected using FCBF algorithm for each TF protein. The features are ordered based on their selection rates (highest to smallest).*

| Dataset | Top 6 Selected Features and their selection rates |
| --- | --- |
| ABF1_YPD | PWM[100], Twist_shift (CC/GG)[90.3], RC[9.4], G content[4.9], Clash(AA/TT)[2.8], Tilt_rise(CC/GG)[1.4] |
| CBF1_SM | Phylo.[99.4], PWM[97.9], Tip(GA/CT)[15.3], Twist_shift (GC/CG)[11.2], Rise (CG/GC)[4.5], CS[2.1] |
| CIN5_YPD | Phylo.[100], PWM[100], Tip(CG/GC)[1.4], RC[0.4], Twist_rise(AG/TC)[0.4], Tip(MEAN)[0.3] |
| DIG1_BUT14 | Phylo.[100], PWM[100], Slide (TA/AT)[78.2], AG content[20.3], Twist_rise(AT/TA)[9.5], Tip(TA/AT)[7.5] |
| FHL1_YPD | PWM[100], Tip(AC/TG)[67.5], G content[21.2], Twist_shift (AA/TT)[19.2], Clash(CC/GG)[9.2], Major Groove (AC/TG)[4.2] |
| FKH1_YPD | Phylo.[100], PWM[46], DNA-BAS(MEAN)[32.6], DNA-BAS(WIDTH)[21.4], RC[11], Major Groove (MEAN)[0.8] |
| GCN4_SM | Phylo.[100], PWM[100], G content[72.2], Bend(CC/GG)[1.3], Flex_slide (CC/GG)[0.5], Flex_shift (CC/GG)[0.4] |
| HAP1_YPD | Phylo.[100], PWM[100], RC[17.3], Tip(CG/GC)[3.1], Bend(MEAN)[0.3], Twist_shift (CC/GG)[0.3] |
| MBP1_H2O2Hi | DNA-BAS(WIDTH)[80.2], Flex_slide (TA/AT)[65.1], Twist_shift (AA/TT)[57.6], Direction(CA/GT)[32.8], PWM[19.3], Melting Temp.(AC/TG)[16.7] |
| NDD1_YPD | PWM[100], Phylo.[100], AG content[39.5], Rise (CC/GG)[17.8], Twist_shift (TA/AT)[13.6], Melting Temp.(CC/GG)[11.5] |
| NRG1_H2O2Hi | PWM[100], Phylo.[100], Twist_rise(AC/TG)[17.1], Stacking(MEAN)[12.5], Rise (CC/GG)[5.7], Slide (CC/GG)[4.1] |
| PHD1_BUT90 | Phylo.[100], PWM[100], Twist_shift (AC/TG)[0.8], Twist_shift (GC/CG)[0.8], Twist_shift (CA/GT)[0.4], Bend(CC/GG)[0.3] |
| RAP1_YPD | PWM[100], Clash(AG/TC)[50.4], Phylo.[42.9], Clash(CA/GT)[42.2], Stacking(AG/TC)[23.5], Clash(CC/GG)[18.6] |
| REB1_H2O2Lo | PWM[69.7], Clash(CG/GC)[39.8], Clash(GC/CG)[34.9], Twist_shift (MEAN)[31.1], DNA-BAS (WIDTH)[30], RC[30] |
| SKN7_H2O2Lo | Phylo.[100], PWM[100], RC[20.3], Clash(AG/TC)[11.6], Rise (CA/GT)[6.8], G content[4] |
| SOK2_BUT14 | Phylo.[100], PWM[100], RC[52.1], Tip(CA/GT)[32.9], NucOcc[2.6], Rise (AA/TT)[0.8] |
| STE12_Alpha | PWM[100], Phylo.[100], Twist_rise(MEAN)[43.6], Stacking(AT/TA)[35.4], Stacking(MEAN)[32.6], Slide (AT/TA)[12] |
| SUT1_YPD | Phylo.[100], PWM[99.3], Clash(CA/GT)[60.2], Major Groove (AC/TG)[13.7], Clash(AC/TG)[9], Tilt_rise(CG/GC)[6.6] |
| SWI4_YPD | PWM[100], Phylo.[100], Melting Temp.(GC/CG)[54.9], Tilt_rise(GA/CT)[52.9], Slide (AT/TA)[22.5], Major Groove (GC/CG)[19.3] |
| SWI5_YPD | PWM[100], Phylo.[97.8], Twist_shift (AA/TT)[49.2], Major Groove (GC/CG)[46.5], Rise (GC/CG)[22.9], Tilt_rise(AT/TA)[12.3] |
| SWI6_YPD | Phylo.[100], PWM[99.9], Flex_shift (AT/TA)[83.3], Twist_shift (MEAN)[49.2], Major Groove (CG/GC)[41.8], Rise (CG/GC)[23.2] |
| UME6_YPD | Phylo.[100], PWM[98.1], G content[4.3], Tip(CC/GG)[4], Inclination(MEAN)[2], Twist_shift (CC/GG)[1.8] |

**Table 4:** *Performance comparison of DNNESCAN against FSCAN and MatInspector.*

| Dataset | Average Performance |  |  |  |  |  | Best Performance |  |  |  |  |  | MatInspector |  |  |
| --- | --- | --- | --- | --- | --- | --- | --- | --- | --- | --- | --- | --- | --- | --- | --- |
|  | DNNESCAN |  |  | FSCAN |  |  | DNNESCAN |  |  | FSCAN |  |  |  |  |  |
|  | P | R | F1 | P | R | F1 | P | R | F1 | P | R | F1 | P | R | F1 |
| ABF1_YPD | 0.74 | 0.96 | 0.84 | 0.69 | 0.83 | 0.75 | 0.75 | 0.97 | 0.85 | 0.7 | 0.85 | 0.76 | 0.67 | 0.95 | 0.79 |
| CBF1_SM | 0.58 | 0.71 | 0.64 | 0.56 | 0.7 | 0.62 | 0.59 | 0.73 | 0.65 | 0.57 | 0.75 | 0.65 | 0.55 | 0.73 | 0.63 |
| CIN5_YPD | 0.3 | 0.75 | 0.43 | 0.21 | 0.59 | 0.31 | 0.34 | 0.74 | 0.47 | 0.23 | 0.57 | 0.33 | 0.2 | 0.64 | 0.3 |
| DIG1_BUT14 | 0.52 | 0.82 | 0.64 | 0.4 | 0.84 | 0.54 | 0.54 | 0.84 | 0.66 | 0.45 | 0.82 | 0.58 | 0.26 | 0.89 | 0.41 |
| FHL1_YPD | 0.33 | 0.66 | 0.44 | 0.38 | 0.72 | 0.49 | 0.36 | 0.69 | 0.48 | 0.4 | 0.74 | 0.52 | 0.29 | 0.92 | 0.44 |
| FKH1_YPD | 0.55 | 0.76 | 0.63 | 0.51 | 0.78 | 0.61 | 0.56 | 0.81 | 0.66 | 0.53 | 0.81 | 0.64 | 0.38 | 0.74 | 0.5 |
| GCN4_SM | 0.57 | 0.75 | 0.65 | 0.5 | 0.67 | 0.57 | 0.57 | 0.79 | 0.66 | 0.51 | 0.73 | 0.6 | 0.49 | 0.76 | 0.6 |
| HAP1_YPD | 0.49 | 0.92 | 0.64 | 0.35 | 0.81 | 0.49 | 0.51 | 0.95 | 0.66 | 0.37 | 0.85 | 0.52 | 0.59 | 0.46 | 0.51 |
| MBP1_H2O2Hi | 0.48 | 0.53 | 0.5 | 0.47 | 0.62 | 0.53 | 0.5 | 0.61 | 0.55 | 0.53 | 0.64 | 0.58 | 0.42 | 0.98 | 0.59 |
| NDD1_YPD | 0.27 | 0.78 | 0.4 | 0.15 | 0.63 | 0.24 | 0.28 | 0.82 | 0.42 | 0.16 | 0.68 | 0.25 | 0.23 | 0.61 | 0.34 |
| NRG1_H2O2Hi | 0.56 | 0.73 | 0.63 | 0.42 | 0.68 | 0.52 | 0.56 | 0.79 | 0.66 | 0.46 | 0.7 | 0.55 | 0.34 | 0.79 | 0.47 |
| PHD1_BUT90 | 0.49 | 0.72 | 0.58 | 0.44 | 0.64 | 0.52 | 0.5 | 0.74 | 0.6 | 0.45 | 0.66 | 0.54 | 0.3 | 0.58 | 0.4 |
| RAP1_YPD | 0.42 | 0.78 | 0.55 | 0.38 | 0.73 | 0.5 | 0.47 | 0.82 | 0.6 | 0.39 | 0.74 | 0.51 | 0.33 | 0.71 | 0.45 |
| REB1_H2O2Lo | 0.82 | 0.92 | 0.87 | 0.82 | 0.89 | 0.85 | 0.83 | 0.93 | 0.88 | 0.83 | 0.9 | 0.86 | 0.84 | 0.9 | 0.87 |
| SKN7_H2O2Lo | 0.32 | 0.88 | 0.47 | 0.22 | 0.65 | 0.33 | 0.35 | 0.92 | 0.51 | 0.23 | 0.69 | 0.35 | 0.35 | 0.46 | 0.39 |
| SOK2_BUT14 | 0.3 | 0.8 | 0.44 | 0.24 | 0.75 | 0.36 | 0.31 | 0.83 | 0.46 | 0.26 | 0.79 | 0.39 | 0.2 | 0.62 | 0.3 |
| STE12_Alpha | 0.39 | 0.71 | 0.51 | 0.3 | 0.75 | 0.43 | 0.41 | 0.74 | 0.53 | 0.32 | 0.76 | 0.45 | 0.31 | 0.47 | 0.37 |
| SUT1_YPD | 0.36 | 0.82 | 0.5 | 0.25 | 0.72 | 0.37 | 0.37 | 0.82 | 0.51 | 0.26 | 0.72 | 0.38 | 0.25 | 0.52 | 0.34 |
| SWI4_YPD | 0.66 | 0.75 | 0.7 | 0.57 | 0.84 | 0.68 | 0.68 | 0.75 | 0.71 | 0.58 | 0.89 | 0.7 | 0.68 | 0.52 | 0.59 |
| SWI5_YPD | 0.36 | 0.6 | 0.45 | 0.3 | 0.69 | 0.41 | 0.41 | 0.62 | 0.49 | 0.33 | 0.67 | 0.44 | 0.27 | 0.7 | 0.39 |
| SWI6_YPD | 0.49 | 0.61 | 0.54 | 0.43 | 0.74 | 0.54 | 0.51 | 0.65 | 0.57 | 0.44 | 0.74 | 0.55 | 0.39 | 0.84 | 0.54 |
| UME6_YPD | 0.78 | 0.88 | 0.83 | 0.74 | 0.86 | 0.8 | 0.79 | 0.89 | 0.84 | 0.77 | 0.88 | 0.82 | 0.7 | 0.95 | 0.81 |
| Average | 0.49 | 0.77 | 0.59 | 0.42 | 0.73 | 0.52 | 0.51 | 0.79 | 0.61 | 0.44 | 0.75 | 0.54 | 0.41 | 0.72 | 0.5 |
| SSR | None |  |  | 20 / 22 |  |  | None |  |  | 20 / 22 |  |  | 21 / 22 |  |  |

### 3.2 Comparison with other methods

In this section, a comparison study is conducted to assess DNNESCAN’s performance against FSCAN (Wang *et al.*, 2016) and the threshold-based system MatInspector (Quandt *et al.*, 1995). There are other approaches for TFBS identification that use machine learning techniques as mentioned in Section ??. However, they use different evaluation protocols to us and different datasets and their source codes are not available to run assessments on our sequence-sets. To appropriately compare the performance of other approaches with DNNESCAN, we report the average and best generalization performance of FSCAN and DNNESCAN over the 10-fold cross-validation procedure that is repeated ten times. On the other hand, we report the performance of MatInspector at the threshold that gives the best F1-measure on the whole sequence-set. It is obvious that the evaluation procedure is in favor of MatInspector and strict for DNNESCAN and FSCAN. It can be seen from the last two rows in Table 4 that DNNESCAN outperforms FSCAN and MatInspector to a large extent. The average positive predictive value of DNNESCAN over all datasets is 49% on average and 51% in the best case while it is 42% and 44% for FSCAN, respectively. The precision of MatInspector is 41% on average. Similarly, it can be observed that the recall of DNNESCAN is 77% on average over all datasets while it is 73% and 72% for FSCAN and MatInspector, respectively, though DNNESCAN can achieve a recall as high as 79% in the best case. Combining the precision and recall performance metrics in the F1-measure shows that DNNESCAN significantly improved the prediction performance of TFBSs by 13% and 18% compared with FSCAN and MatInspector, respectively. Furthermore, DNNESCAN increased the F1-measure as high as 67% and 56% for some TF proteins compared with FSCAN and MatInspector, respectively (see DIG1 and NDD1 in Table 4).

By comparing the three methods on individual sequence-sets, we can observe that DNNESCAN outperforms FSCAN and MatInspector on 20 and 21, out of the 22 datasets, respectively. The performance of DNNESCAN falls by 15% compared with MatInspector on one dataset only (MBP1) due to the low recall rate. It can also be seen from Table 4 that the prediction of TFBSs is boosted by at least 20% for eleven TFs if we compare the average F1-measure of DNNESCAN with that of MatInspector on CIN5, DIG1, FKH1, HAP1, NRG1, PHD1, RAP1, SKN7, SOK2, STE12 and SUT1. DNNESCAN was able to drastically reduce the number of false positives in most of the cases due to the large amount of prior knowledge that it uses to characterize TFBSs. Interestingly, an 8% drop in the recall of DNNESCAN on the DIG1 and NRG1 datasets resulted in 100% and 65% gain, respectively, on the precision of our system compared with MatInspector. On the contrary, when the precision of DNNESCAN falls by 17% and 9% on the HAP1 and SKN7 datasets, respectively, its recall jumps by 100% and 91%, respectively, compared with MatInspector. This behaviour in our system could be attributed to the relevance of the K-mer features to the investigated TF protein. Therefore, the DNNE learning algorithm that forms the classification boundary between background K-mers and functional binding sites plays a key role in balancing the trade-off between precision and recall. Overall, DNNESCAN convincingly outperforms the threshold-based approach in 18 cases of the 22 studied ChIP-chip datasets and performs comparably on three datasets. Similar observations can be drawn when DNNESCAN is compared with FSCAN though there is no significant performance improvement on many datasets as with MatInspector. Table 4 shows a gain of more than 20% on the DNNESCAN F1-measure for seven TFs, namely CIN5, HAP1, NDD1, NRG1, SKN7, SOK2 and SUT1. This improvement results from an increase in both precision and recall on these datasets. For example, the precision and recall of DNNESCAN rose by 80% and 24%, respectively, on the NDD1 dataset. These results affirm the role of prior knowledge in identifying TFBSs.

### 3.3 Scanning for novel binding sites

In this section, transcription factor DNNESCAN models are created for the 22 TFs without partitioning sequence-sets and the models that maximize F1-measure are selected. Then, the bound sequence-set of TFs are scanned to identify new potential binding sites in these intergenic regions (IGRs). To verify the newly discovered binding sites, transcriptional regulatory evidence for the yeast genome is searched in the literature. Our verification strategy is based on the assumption that if (i) a given gene is known to be regulated by the investigated TF protein, (ii) this regulatory relationship has been *verified* by evidence different from the ChIP-chip experiments of (MacIsaac *et al.*, 2006), (iii) a potential binding site is detected by DNNESCAN upstream this target gene, and (iv) there is no known binding site for the investigated TF in the IGR upstream this target gene, then the detected binding site is more likely to be a *novel* binding site on which the investigated TF could bind to regulate the target gene. In order to implement this verification strategy, the experimentally *verified* target genes of each investigated protein were collected from SGD database on 03 March 2014 (Cherry *et al.*, 2012). Next, some of the novel binding sites for the 22 studied TFs that are reported by DNNESCAN are discussed.

Two binding sites **atcattacacacg** and **gtcactacaaacg** are found for ABF1 in the IGR between PEX3 and UBX5 at positions 1127702 and 1127819, respectively, of chromosome IV. It is well-known in the literature using microarray experiments that ABF1 regulates PEX3 (Yarragudi *et al.*, 2007) and the discovered TFBSs are within 300bp from the transcription start site (TSS) of PEX3. Two putative binding sites **gtcacgtg** and **atcacgtg** are located for CBF1 at positions 289750 and 289752 of chromosome XII, respectively, and upstream GAL2 which is regulated by CBF1. It is evident that the removal of CBF1 leads to changes in the chromatin structure at the GAL2 promoter (Mellor *et al.*, 1990). Two of the CIN5 binding sites **tttacctaa** and **atcacctaa** are recognized in chromosome XV between SPS4 and SFG1 and it was verified by several experiments that CIN5 is a regulator to SFG1 (Tan *et al.*, 2008; Venters and Pugh, 2009). One of the sites is within 100bp from the TSS of SFG1. Four TFBSs for DIG1 occur in the region between RNQ1 and FUS1 and it was confirmed using RNA expression microarrays that FUS1 is regulated by DIG1 (Hu *et al.*, 2007). These sites are located within 200 bp from the TSS of FUS1 that is involved in the mating and growth pathway (Cherry *et al.*, 2012). Furthermore, DIG1 is involved in the regulation of mating-specific genes and the invasive growth pathway of yeast (Cook *et al.*, 1996). As for the FHL1 TF, it binds the DNA directly at highly active ribosomal protein genes (Hermann-Le Denmat *et al.*, 1994). One of its discovered BSs is **atgcacgggt** which is detected at position 555512 of chromosome VII and upstream two genes TIM21 and RPL26B that are evidently regulated by FHL1 (Schawalder *et al.*, 2004). It can be noted that this regulatory sequence is closer to the TSS of RPL26B which is a component of the large ribosomal subunit (60S) (Nakao *et al.*, 2004). FKH1 is involved in the transcriptional regulation of several genes during the G2/M phase (Hollenhorst *et al.*, 2000). Two binding sites **gtaaacag** and **gtaaacaa** are predicted for FKH1 in chromosome VIII upstream DSE2 that is involved in the mitotic cell cycle and known to be regulated by FKH1 (Shapira *et al.*, 2004). Furthermore, one binding site **atgactct** is found for GCN4 at position 449456 upstream YMC2 which is evidently regulated by GCN4 according to several microarray experiments (Staschke *et al.*, 2010).

In addition to this, a regulatory element for HAP1 is detected upstream HAP1 itself at position 646024 of chromosome XII. Interestingly, electrophoretic mobility shift assay (EMSA) shows that HAP1 binds directly to the DNA and down-regulates the expression of its own gene (Xin *et al.*, 2007). Indeed, (Xin *et al.*, 2007) reported that HAP1 binds its own promoter within −341 to −380 from the TSS and our discovered TFBS is within this genomic region. MBP1 regulates DNA synthesis and repair genes by binding to the regulatory element MLuI-box in their promoters (Koch *et al.*, 1993). Two MBP1 potential binding sites of the same sequence **gcgcgtc** are detected at different positions upstream CLB5 and within 400 bp from its TSS. CLB5 is involved in DNA replication during S phase, activates Cdc28p to promote the initiation of DNA synthesis and functions in formation of mitotic spindles (Schwob and Nasmyth, 1993). It has been found in the literature that MBP1 is an important expression regulator for CLB5 (Bean *et al.*, 2004). NDD1 regulates around 107 target genes in the yeast genome, but due to a lack of experiments on NDD1, we cannot verify any of the discovered binding sites. Two NRG1 binding sites **ggaccct** and **agaccct** are found in chromosome XIII at positions 915090 and 915097, respectively, and upstream the plasma membrane transporter gene FET4. It has been reported in the literature that NRG1 is a regulator for FET4 (Goh *et al.*, 2010) and the detected TFBS is within 600 bp from FET4. Six potential sites for PHD1 are found in chromosome X between IME1 and RPL43B in the 2653bp upstream region of both of them. ChIP-exo (exonucleases) experiments conducted by (Rhee and Pugh, 2011) showed that PHD1 binds on four DNA elements in this intergenic region. One of these elements overlaps with the site discovered by DNNESCAN **gggcac** at position 606763. Five RAP1 binding sites are found in chromosome II upstream genes of three ribosomal proteins (RPL19A, RPL19B and RPS11B) that are evidently regulated by RAP1 (Hu *et al.*, 2007). Actually, three of these regulatory sites (**acccaaacat** at position 332543 and upstream RPS11B, **atccagacat** at position 415668 and upstream RPL19A, and **acccatgcat** at position 415684 and upstream RPL19A) overlap with RAP1 bound elements that are found by (Rhee and Pugh, 2011) using ChIP-exo experiments. Interestingly, 19 of the 20 TFBSs discovered for REB1 occur in REB1 bound regions as reported by ChIP-exo experiments (Rhee and Pugh, 2011) and one binding site is in a close proximity from a REB1 bound region.

SKN7 regulates genes that are involved in the cell wall biosynthesis, the cell cycle and the oxidative and osmotic stresses response (Fassler and West, 2011). An important SKN7 binding site discovered by DNNESCAN is **gccggccg** at position 707247 of chromosome II and upstream GPX2 whose expression is regulated by SKN7 according to microarray assays (Kelley and Ideker, 2009). (Tsuzi *et al.*, 2004) reported that SKN7 is essential for the oxidative stress response of GPX2 and it binds to its promoter sequence. Moreover, SOK2 mainly regulates genes involved in filamentous growth and cell wall adhesion (Ward and Garrett, 1994). Five separated TFBSs for SOK2 are hit by DNNESCAN in the IGR between MTC2 and YKL096C-B upstream CWP2 which is reported to be regulated by SOK2 (Chua *et al.*, 2006). CWP2 is a major constituent of the cell well in the form of covalently linked mannoprotein and plays a key role in stabilizing the cell wall (Cherry *et al.*, 2012). STE12 activates genes involved in mating or pseudohyphal growth pathways (Bardwell *et al.*, 1998). Four STE12 binding sites **tgagaca**, **tgaaaca**, **tgaaacg** and **tgaaaca** are detected in chromosome III upstream FUS1 and within 200 bp from its TSS. It has been found that FUS1 is a membrane protein which is required for cell fusion and its expression seems to be regulated by the mating pheromone (Cherry *et al.*, 2012). Microarray assays also confirm that the expression of FUS1 is regulated by the binding of STE12 on its promoter regardless of the existence of the TEC1 protein (Heise *et al.*, 2010). DNNESCAN detected 302 potential binding sites for SUT1 in the SUT1_YPD sequence-set. However, we could not verify any of these sites using gene transcriptional regulatory networks. SWI4 requires SWI6 in order to bind to DNA regulatory elements since it makes a complex binding factor (SBF) with it which induces DNA conformational changes that allow SWI4 to access DNA bases. SWI4 mainly regulates genes involved in the G1/S cell cycle including G1 cyclins and the HO endonuclease (Stern *et al.*, 1984). A binding site **cgcgaaa** for SWI4 is hit by DNNESCAN upstream PLM2 at 206bp from its TSS. (Bean *et al.*, 2005)’s microarray experiments demonstrate that the expression of PLM2 is controlled by SWI4 and PLM2 is a target for the SBF transcription complex. On the other hand, SWI5 regulates genes involved in cell separation and mating type switching in daughter cells (Stern *et al.*, 1984). Five SWI5 binding sites are detected in chromosome X upstream PRY3 within 400 bp from its TSS. Experimental results show that the expression of PRY3 is regulated by SWI5 (Di Talia *et al.*, 2009) and it has been found that PRY3 plays an important role in mating efficiency (Cherry *et al.*, 2012). Interestingly, one of the detected sites is at 49 bp from PRY3 ORF while the others occur at different positions in its promoter. As mentioned earlier, SWI6 forms the transcriptional complex SBF along with SWI4, yet it forms the complex MBF with MBP1. As a result, it regulates genes at the G1/S transition, G1 cyclins and genes involved in DNA synthesis and repair (Breeden and Nasmyth, 1987). Two regulatory elements for SWI6 are hit upstream TOS6 at positions 65309 (**gacgcg**) and 65342 (**gacgcg**) of chromosome XIV within 500 bp from its TSS. Actually, microarray RNA expression assays in a rich medium where SWI6 is removed show that SWI6 is important for TOS6 to be transcribed (Hu *et al.*, 2007). Finally, UME6 regulates genes involved in meiosis and sporulation, and functions as a repressor for several early meiosis-specific genes during vegetative growth (?). Microarray assays in a rich medium growth condition showed that the expression of ULP2 and BCY1 in chromosome VIII could be regulated by UME6 (Hu *et al.*, 2007). Two binding sites of the same DNA sequence **tagccgccga** were discovered by DNNESCAN in the 963 bp IGR between these two genes at positions 292143 and 292214. Furthermore, the meiosis-specific telomere protein NDJ1 in chromosome XV was found to be regulated by UME6 in (Hu *et al.*, 2007)’s assays. A potential UME6 binding site **tagccgccca** is detected by DNNESCAN within 80 bp from NDJ1. For the complete list of verified TFBSs discovered by DNNESCAN, see the supplementary files.

## 4 CONCLUSION

In this manuscript, we proposed a computational intelligence framework for TFBS identification. Our proposed system, named DNNESCAN, is a threshold-free system and uses a recent ensemble learning algorithm called DNNE to create classifier models for TFBSs. Unlike other TFBS predictors published in the literature (Hooghe *et al.*, 2012; Bauer *et al.*, 2010; Meysman *et al.*, 2011; Steffen *et al.*, 2002; Pudimat *et al.*, 2005), DNNESCAN uses systematic preprocessing procedures in order to prepare training and testing datasets. Moreover, the data imbalance reduction technique was used to alleviate the highly imbalance ratio between known binding sites and background K-mers in the training datasets. In order to better characterize TFBSs, prior knowledge composed of motif-dependent, conformational, thermodynamics, phylogenetic and letter-based features is utilized along with a feature selection algorithm that helps reduce classifier complexity. For the purposes of this research, DNNESCAN models are generated for 22 TF proteins from the S. Cerevisiae genome. These models can be used to scan DNA sequences and locate putative binding sites. Although ChIP-chip sequence-sets are used in our experiments, the proposed framework can be easily extended to any set of co-regulated promoter sequences.

The simulation results show that DNNESCAN significantly outperforms threshold-based systems like MatInspector (Quandt *et al.*, 1995) and reduces the number of false binding sites effectively. The features selected by DNNESCAN to characterize the binding sites of individual TFs provide new insights on the binding specificities of these TFs. Finally, DNNESCAN was able to discover new binding sites that could be verified using experimental studies published in the literature. This highlights the efficiency and efficacy of our proposed framework. Further work can be extended on this framework so that a post-processing procedure is implemented to increase prediction accuracy. New prior knowledge on the protein-DNA binding specificity can be easily integrated in the DNNESCAN framework and hence better performance results could be obtained.

## Acknowledgment

M. A. was receiveing a PhD scholarship from La Trobe University and a Top-Up Ph.D. scholarship from the Victorian Life Sciences Computation Initiative (VLSCI) during the development of this research.

